# Binding partner regulation of Myosin VI: Loss of tumour-suppressor Dab2 leads to enhanced activity of nuclear myosin

**DOI:** 10.1101/639963

**Authors:** Natalia Fili, Yukti Hari-Gupta, Bjork Aston, Ália dos Santos, Rosemarie E. Gough, Bana Alamad, Lin Wang, Marisa Martin-Fernandez, Christopher P. Toseland

## Abstract

Myosin VI is involved in a variety of cellular processes ranging from endocytosis to transcription. This multi-functional potential is achieved through alternative isoform splicing and through the interaction with a diverse network of binding partners. However, the interplay between the two modes of regulation remains unexplored. To this end, we have compared two different binding partners, Dab2 and CALCOCO2/NDP52, and their interaction with two myosin VI splice isoforms. We found that both isoforms adopt an auto-inhibited state and are subsequently activated by binding partner association. However, differential regulation is achieved through a high and a low affinity binding motifs within myosin VI, with one isoform having the high affinity site blocked. This allows competition between partners and links isoform splicing with binding partner selectivity. Dab2 competition hinders the activity of nuclear myosin VI by preventing DNA binding and transcription. Moreover, re-introduction of Dab2 in the Dab2-deficient MCF-7 cells leads to a decrease in myosin VI-dependent estrogen receptor gene expression. We propose that the frequent loss of Dab2 during the onset of cancer enables a higher level of nuclear myosin VI activity, thereby driving the activity of the estrogen receptor to promote tumourgenesis.

## Introduction

Myosin VI (MVI) is an actin-based molecular motor which performs numerous vital roles in key cellular processes such as cell migration, endocytosis, exocytosis and transcription (Fili et al., 2017; Roberts et al., 2004; Vreugde et al., 2006). Defects in MVI lead to various diseases including hypertrophic cardiomyopathy, deafness and cancer (Avraham et al., 1995; Dunn et al., 2006; Mohiddin et al., 2004; Yoshida et al., 2004). MVI consists of the highly conserved actin-binding motor domain, a neck region and a C-terminal globular cargo binding domain (CBD) (Figure 1a). We have recently shown that MVI can adopt a back-folded conformation, in which the CBD is brought in close proximity to the motor domain (Fili et al., 2017). Moreover, two regions within the tail (MVI_TAIL_, aa 814-1253) can be alternatively spliced, resulting in a 31 residue insertion (large-insert, LI) adjacent to the CBD, and/or an 9 residue insertion in the middle of the CBD (small-insert, SI) (Buss et al., 2001). This leads to several splice isoforms, namely the non-insert (NI), SI, LI and LI+SI, each with distinct intracellular distributions and functions (Au et al., 2007; Buss et al., 2001). For example, the NI isoform is able to enter the nucleus, whereas the LI is confined to the cell periphery (Fili et al., 2017). The intracellular localisation and function of MVI is also regulated through its interaction with a broad range of binding partners, such as disabled-2 (Dab2), the GAIP-interacting protein C-terminus (GIPC) and the nuclear dot protein 52 (NDP52). These partners specifically bind to one of two established motifs within the CBD of MVI, namely the RRL and WWY (Morriswood et al., 2007; Naccache et al., 2006; Spudich et al., 2007). NDP52/CALCOCO2 is an RRL binding partner of MVI. It was initially identified in the nucleus (Korioth et al., 1995), but it was later found to be mostly cytoplasmic (Sternsdorf et al., 1997), with roles in cell adhesion and autophagy (Morriswood et al., 2007; Mostowy et al., 2011). NDP52 has been shown to release the back-folded conformation of MVI, allowing MVI to dimerize and to interact with DNA, both of which enable coupling of MVI to RNA Polymerase II. Moreover, NDP52 has been shown to have a role in regulating transcription (Fili et al., 2017), possibly as a co-activator, similarly to its highly conserved family member CoCoA (Yang et al., 2006). Dab2, also known as Differentially expressed in Ovarian Carcinoma (DOC-2), links MVI to clathrin-coated vesicles at the early stages of endocytosis (Spudich et al., 2007), through interaction with the WWY motif. Dab2 is down-regulated in majority of breast and ovarian cancers. Moreover, depletion and re-expression of Dab2 can trigger tumorigenesis or suppress growth, respectively (He et al., 2001). Therefore, Dab2 is considered as a tumour suppressor. The selectivity of MVI for its binding partners is in part regulated by isoform splicing. The LI encodes an alpha helix which sits upon, and therefore blocks, the RRL motif (Wollscheid et al., 2016). This prevents partners, such as NDP52, interacting with the protein, and therefore the binding partner interactions of this isoform are driven by the WWY motif. In contrast, in the NI isoform, both the RRL and WWY motifs are available for binding. In this case, binding partner selectivity would be an important regulatory mechanism. In the light of our recent work on the regulation of the NI isoform by NDP52 (Fili et al., 2017) and with the aim to unravel such a regulatory the mechanism, we have first investigated biochemically whether interactions with the WWY motif can similarly regulate both back-folding and dimerization of MVI, in both the NI and LI isoforms. Moreover, we have established how competition between the two binding sites is achieved and how this can impact upon the role of nuclear MVI in gene expression.

**Fig. 1.**
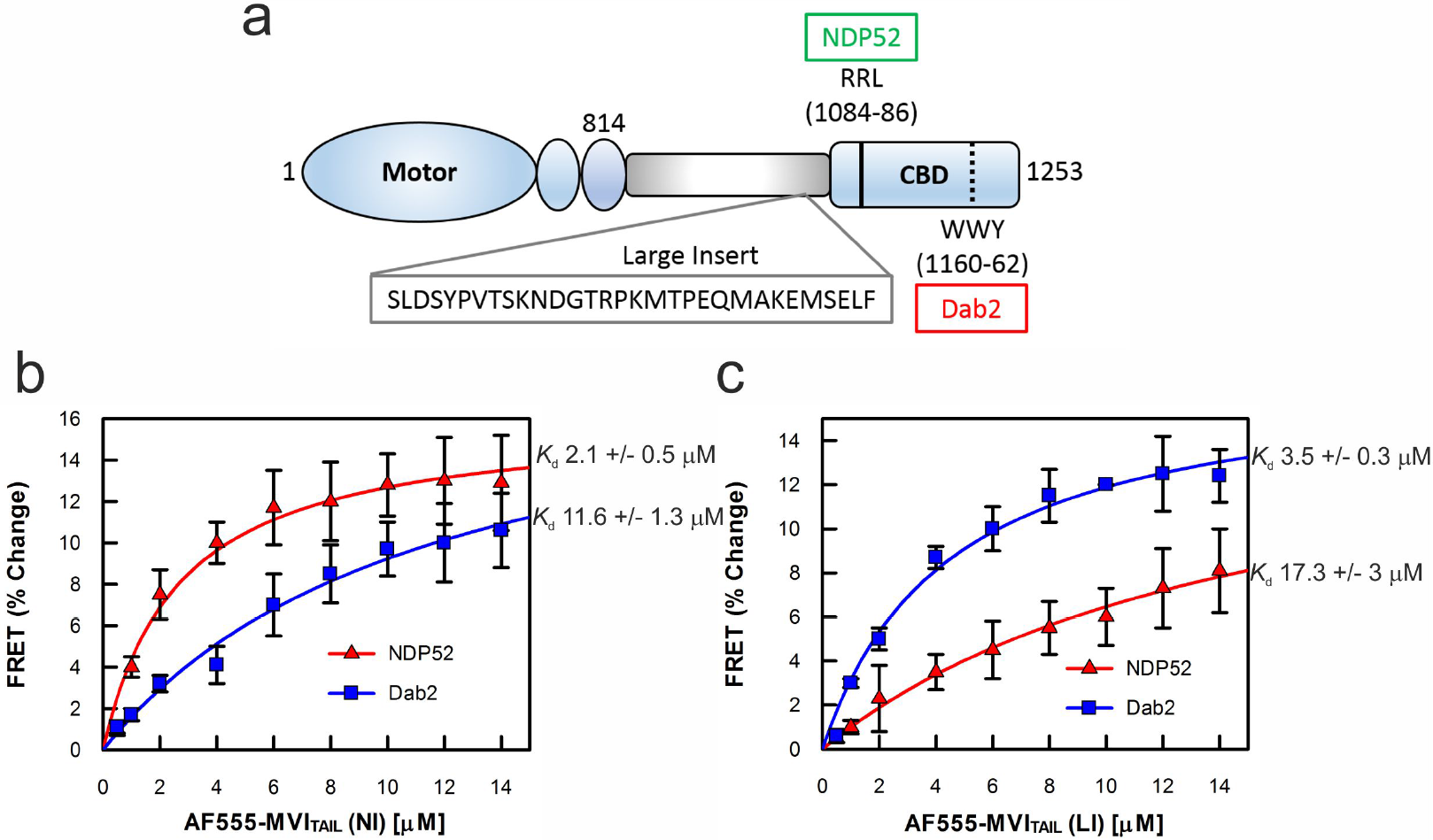
Interaction between Myosin VI and binding partners NDP52 and Dab2. (a) Cartoon depiction of the key regions of the MVI_TAIL_, as discussed in the text. (b) FRET titration of MVI_TAIL(NI)_ against 1 μM NDP52 (red triangles) or Dab2 (blue squares). (c) FRET titration of MVI_TAIL(LI)_ against 1 μM NDP52 (red triangles) or Dab2 (blue squares). All titration data fitting was performed as described in Methods giving a (*K*_d_ as plotted (Error bars represent SEM from three independent experiments).

## Results

### Interactions between binding partners and myosin VI

In order to establish how the selectivity of MVI for its binding partners is regulated, we compared its interactions with two binding partners, namely NDP52 and Dab2, as representatives of RRL and WWY binding proteins, respectively. Given our recent work on the interaction of the NI isoform with NDP52 (Fili et al., 2017), here we focus on the interaction with Dab2. Before investigating the effect of Dab2 upon MVI, we first assessed their interaction. Recombinant full length Dab2 was highly unstable and unable to yield sufficient amounts of protein for biochemical characterisation. Therefore, we used the stable C-Terminal region of the protein (residues 649-770), which contains the MVI binding site (Spudich et al., 2007). This truncation of Dab2 will be referred to as Dab2 throughout the manuscript, unless stated. To characterise the interaction between MVI and Dab2, we performed an in vitro FRET assay by titrating Alexa555-MVI_TAIL(NI)_ or Alexa555-MVI_TAIL(LI)_ against FITC-Dab2. As demonstrated by the binding curves in Figures 1b and 1c, Dab2 displayed relatively weak binding to the MVI_TAIL(NI)_ (*K*_d_ 11.6 μM). Binding was noticeably enhanced for the MVI_TAIL(LI)_ (*K*_d_ 3.5 μM), suggesting that the LI stabilises the interaction with Dab2. For comparison, measurements were also performed with NDP52. Consistent with the previous results (Fili et al., 2017), MVI_TAIL(NI)_ bound to NDP52 with a low micromolar affinity (*K*_d_ 2.1 μM). In contrast, binding to the MVI_TAIL(LI)_ was over 8-fold weaker (*K*_d_ 17.3 μM), indicating that NDP52 selectively interacts with MVI_TAIL(NI)_, rather than the MVI_TAIL(LI)_. Based on this data, the differential affinity between the WWY and RRL sites suggests that the NI isoform would selectively bind RRL binding partners over its WWY competitors. In contrast, the LI isoform shows preferential binding to the WWY partners because the LI helix (i) masks the higher affinity RRL motif (Wollscheid et al., 2016), thereby impeding NDP52 binding, and (ii) may provide additional interaction sites to increase the Dab2 affinity. To further explore these interactions with full length proteins, we assessed their association within cells. In HeLa cells, which only express the NI isoform (Puri et al., 2010), endogenous MVI shows little co-localisation with endogenous Dab2, if any (Figure 2a). In contrast, transiently expressed GFP-LI showed significant co-localisation with endogenous Dab2, as highlighted by the 10-fold increase in Pearson’s coefficient (Figure 2b and 2d). Consistent with the titration measurements, these observations support that the selectivity of MVI for its partners differs depending on the isoform. Our biochemical data suggested that the association of MVI with its partners is a concentration dependent process. To address this in the cellular environment, we artificially increased the intracellular levels of the NI isoform by transient overexpression of GFP-NI-MVI and assessed its colocalization with endogenous Dab2. Consistent with our titration data, increase in the NI intracellular levels shifted the extent of colocalization between the two proteins at levels comparable to the LI-MVI (Figure 2b and 2d). To confirm the specificity of this observation, we also assessed the effect of transiently over-expressing two mutants of the NI isoform, each carrying mutations that abolish one of the two binding motifs (Figure 2c and 2d). As expected, the GFP-NI-MVI (WWY/WLY), in which the WWY binding site is abolished, did not show any colocalization with Dab2. In contrast, mutation of the RRL motif did not affect the colocalization with Dab2. In fact, there was a slight increase in colocalization, which may relate to an increase in free MVI available to interact with WWY binding partners. Overall, these observations are consistent with the conclusions from the titration measurements which suggest that significant interactions between the NI MVI_TAIL_ and Dab2 can occur only at high protein concentrations. Therefore, our data support that the WWY binding site has a weaker affinity for protein-protein interactions.

**Fig. 2.**
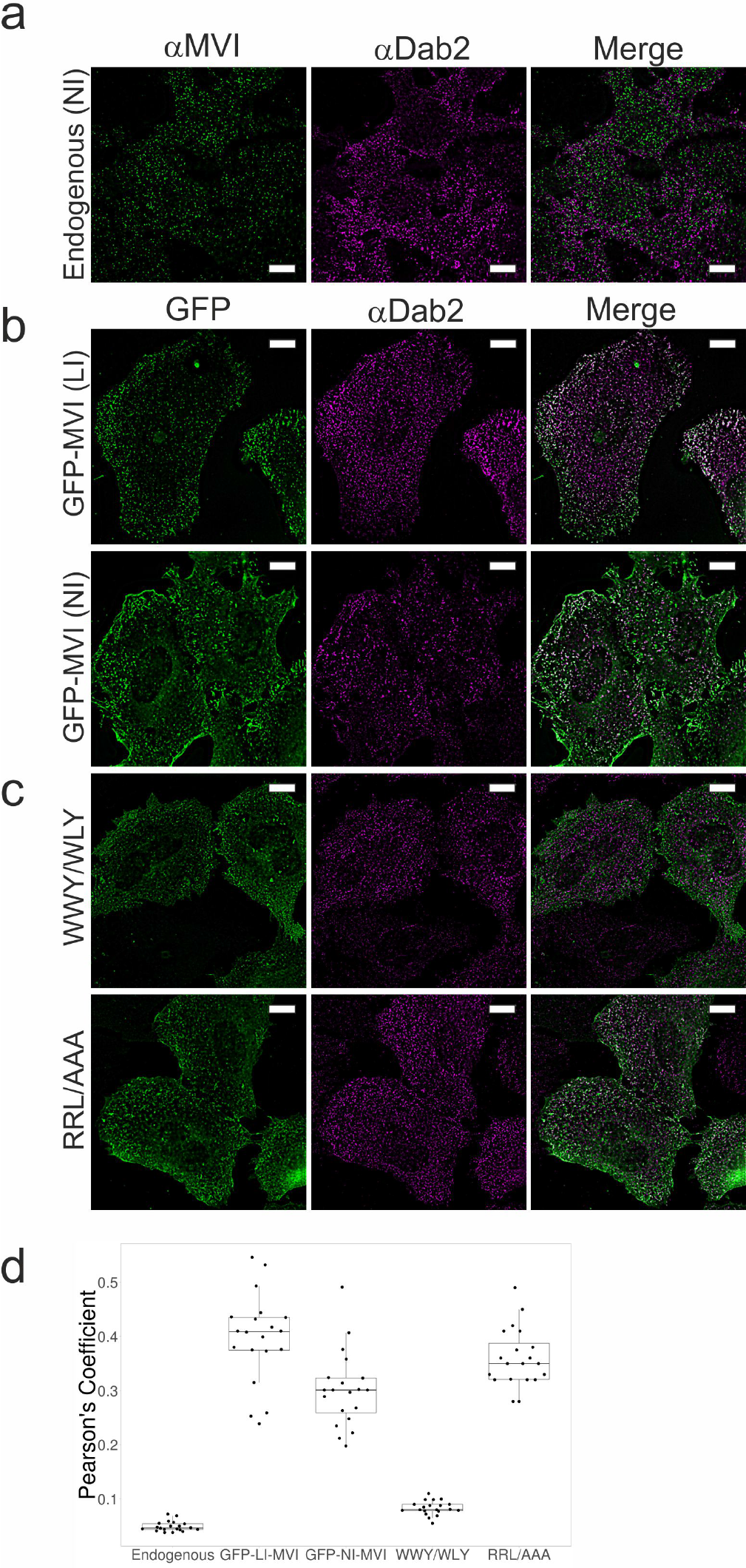
Myosin VI interaction with Dab2 in HeLa cells. (a) Immunofluorescence staining against MVI (green) and Dab2 (magenta) in HeLa cells. (b) Representative images of transiently expressed NI- and LI-GFP-MVI in HeLa cells combined with immunofluorescence staining against Dab2. (c) Representative images of transiently expressed NI-MVI mutants WWY/WLY and RRL/AAA in HeLa cells combined with immunofluorescence staining against Dab2. Scale bar 10 μm in all images. (d) Pearson’s coefficient for MVI colocalisation with Dab2 from images in a-c. Figure was generated using (Postma and Goedhart, 2019)

### The large insert isoform is back-folded

We have previously revealed that the NI-MVI isoform is back-folded in vitro and in cells (Fili et al., 2017). To address whether back-folding is general feature of MVI that does not depend on the isoform, we investigated the conformation of LI-MVI. We utilised a FRET-based assay by titrating Alexa555-CBD against FITC-_N_MVI_TAIL(LI)_ to assess whether there is an interaction between the N- and C-terminal tail domains. A significant change in FRET was detected, indicating that the two domains are in close proximity (Figure 3a). The same was observed for the NI isoform (Figure 3b), as previously observed (Fili et al., 2017). To address whether Dab2 can regulate the interaction between the tail domains in the same manner as NDP52, the FRET assay between the _N_MVI_TAIL(NI)_, or _N_MVI_TAIL(LI)_, and the CBD was repeated following pre-incubation of the CBD with an excess of Dab2. As with NDP52, Dab2 sequestered the CBD, preventing the interaction between the two domains (Figures 3a and 3b). However, Dab2 was not as efficient as NDP52, which is consistent with the weaker affinity of Dab2 for MVI. While NDP52 is usually impeded from binding to the LI isoform, here NDP52 could bind to the CBD, as the LI helix was present on the N-terminal tail fragment. To investigate the effect of NDP52 upon LI-MVI unfolding, we used a FRET-based MVI tail conformation reporter, we have previously developed (Fili et al., 2017). Our reporter was based on a GFP-RFP FRET pair with the MVI_TAIL(NI)_ or MVI_TAIL(LI)_, placed in the middle (Figure 3c). As shown by the fluorescent spectra, a high FRET population was observed with both of these reporters, supporting the conclusion that both tails have the ability to backfold. As seen previously (Fili et al., 2017), addition of 5 μM NDP52 to the MVI_TAIL(NI)_ results in loss of the high FRET population. However, addition of NDP52 had little effect, if any, on the MVI_TAIL(LI)_, as expected given their low affinity. In contrast, 5 μM Dab2 was able to deplete the FRET population of the MVI_TAIL(LI)_. However, 20 μM of Dab2 were required to induce an equivalent effect on MVI_TAIL(NI)_, given its low affinity for this tail. This observation highlights both isoform selectivity for binding partner as well as the weaker association between Dab2 and MVI. Taken together, these data demonstrate that intramolecular backfolding is not iso-form specific, but rather an intrinsic feature of MVI. They also highlight how selectivity of MVI for its binding partners and the affinity of these interactions can also regulate the conformation of the protein.

**Fig. 3.**
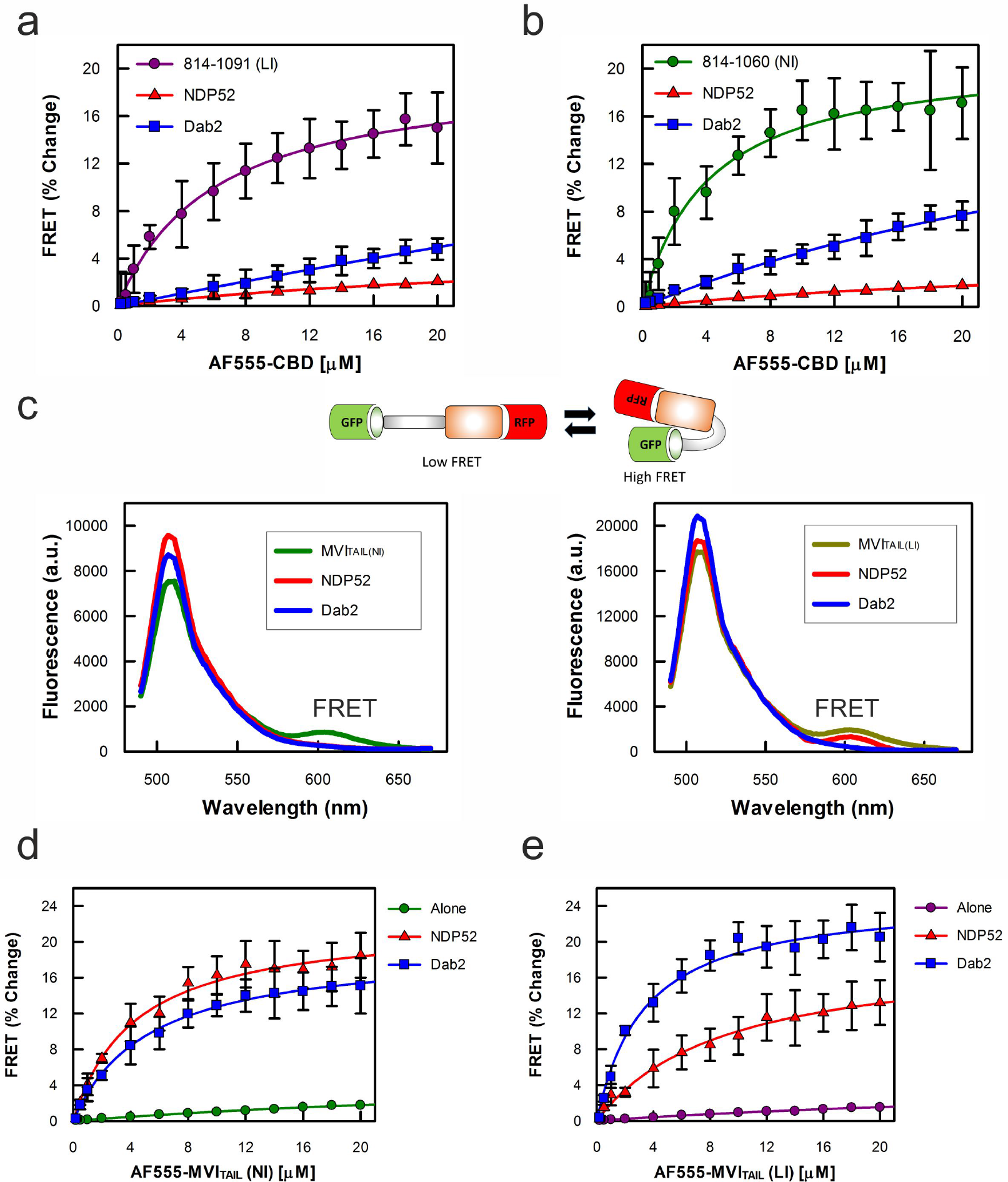
Isoform independent backfolding of myosin VI. (a) FRET titration of CBD against _N_MVI_TAIL(LI)_ (1 μM) in the presence of NDP52 (*K*_d_ N.D.) or Dab2 (*K*_d_ 6 μM), at 10 μM. (b) FRET titration of CBD against _N_MVI_TAIL(NI)_ (1 μM) in the presence of NDP52 (*K*_d_ 4.1 μM) or Dab2 (*K*_d_ N.D.), at 10 μM. (c) Representative fluorescence spectra of (1 μM) GFP-MVI_TAIL(NI)_-RFP +/− 5 μM NDP52, or 20 μM Dab2 (left) and GFP-MVI_TAIL(LI)_-RFP (right) +/− 5 μM NDP52, or Dab2. (d) FRET titration of AF555-MVI_TAIL(NI)_ against 1 μM FITC-MVI_TAIL(NI)_ +/− Dab2 (20 μM) and NDP52 (20 μM). Data fitting generated NDP52 *K*_d_^DIMER^ as 4.3 μM and Dab2 *K*_d_^DIMER^ as 5.3 μM. (e) FRET titration as above but with MVI_TAIL(LI)_. Data fitting generated NDP52 *K*_d_^DIMER^ as 9.3 μM and Dab2 *K*_d_^DIMER^ as 3.3 μM. All titration data fitting was performed as described in Methods (Error bars represent SEM from three independent experiments).

### Dab2 mediates Myosin VI Large Insert isoform dimerization

We have previously revealed how NDP52 interacts with MVI NI to bring about its unfolding (Fili et al., 2017). Unfolding subsequently exposes dimerization sites, leadino protein oligomerization. However, the ability of the LI isoform to dimerise has not been yet explored. The presence of the additional alpha helix in this isoform may perturb its dimerization. Having shown that Dab2 is capable to unfold both NI and LI MVI, we therefore explored whether Dab2 can also lead to oligomerization of both isoforms. To this end, we performed a FRET assay, whereby two pools of MVI_TAIL(NI)_ or MVI_TAIL(LI)_ were labelled, one with FITC and the other one with Alexa555. Titrations revealed a weak association between the two tail pools (Figures 3d and 3e), consistent with our previous results (Fili et al., 2017). Upon addition of 20 μM excess Dab2, a change in FRET signal was observed, indicating the formation of a dimer complex. This occurred with both the NI and LI tails, with the LI signal being higher possibly due the higher association with Dab2. 20 μM excess of NDP52 was able to trigger dimerization of MVI-TAIL(NI) but failed to significantly dimerize the MVI_TAIL(LI)_, consistent with the poor binding of NDP52 to the LI isoform. To further confirm the effect of Dab2 on MVI dimerisation, the steady-state ATPase kinetics of full length MVI-LI was recorded. Following addition of 20 μM Dab2, a decrease in actin-activated ATPase rate from 2.9 s^−1^ to 1.6 s^−1^ was observed. A similar impact was observed with the MVI-NI, with the ATPase rate being decreased from 3.3 s^−1^ to 1.9 s^−1^. This reduction in the rate constant is typical of molecular gating, whereby each head of the dimer alternates ATP hydrolysis (Sweeney et al., 2007). Taking all our data together, we propose the following model: NDP52 specifically associates with MVI-NI through the RRL motif to trigger unfolding of the protein and subsequent dimerization. Similarly, binding of Dab2 to the LI isoform through the WWY motif also leads to MVI unfolding and dimerisation. Whilst Dab2 has also the ability to bind the NI isoform, its weaker affinity for the WWY motif makes this interaction less favourable compared to the high affinity binding of NDP52 to the RRL motif. In this way, the unfolding and dimerization of the NI isoform is preferentially assigned to the RRL binding partners.

### Effect of binding partner competition upon the biochemical properties of nuclear myosin VI

We have shown that association of a binding partner with either the RRL or the WWY motif can bring about similar structural changes within MVI, triggering both its unfolding and dimerization. We have also shown that the selectivity of MVI for its binding partners is regulated by the differential affinity between the two motifs. However, what is the biological impact of this binding partner selectivity and what would be the effect if its gets disrupted? More specifically, what is the effect of Dab2 upon the function of nuclear MVI, which we previously identified as the NI isoform? We have established that Dab2 can interact with this isoform at high protein concentrations. Interestingly, Dab2 can also be detected in the nucleus (Figure 4a) and therefore it could interact with nuclear MVI. We have previously shown that MVI contains a DNA binding site on two conserved loops in the CBD (Supplementary Figure 1a) (Fili et al., 2017). These sites are only exposed upon unfolding of the tail domain and therefore, DNA binding is dependent upon the interaction with a binding partner (Fili et al., 2017). First, we wanted to assess the effect of binding partners upon the ability of MVI to bind DNA, which is critical for its role in transcription (Fili et al., 2017). To determine DNA binding, we measured the fluorescence anisotropy of labelled DNA. As reported previously (Fili et al., 2017), NDP52 independently binds to DNA itself and therefore, it is unsuitable for this purpose. In contrast, Dab2 does not bind DNA (Supplementary Figure 1b) and therefore it can be used to determine its impact upon MVI binding DNA. To this end, we monitored the binding of MVI CBD to fluorescently labelled DNA (Figure 4b). Whereas the CBD alone could bind to DNA with strong affinity (*K*_d_ 140 nM), the presence of Dab2 inhibited DNA binding in a concentration-dependent manner. Interestingly, the DNA binding sites of MVI are in close proximity to the WWY motif where Dab2 binds (Supplementary Figure 1a). Therefore, the interaction of Dab2 with MVI might induce a steric hindrance, or a structural change within the CBD, which prevents the complex from binding DNA. The ability of MVI to bind DNA is important for efficient *in vitro* transcription (Fili et al). Therefore, to assess the effect of Dab2 on the transcription activity of MVI, we performed *in vitro* transcription assays using the HeLaScribe nuclear extracts (Figure 4c). Antibody-depletion of MVI leads to a 70 percent decrease in transcription yield. A similar impact is achieved through the addition of recombinant CBD to displace MVI. Interestingly, addition of 10 μM recombinant Dab2 leads to a 60 percent decrease in transcription. We therefore propose that this effect is due to Dab2 interfering with the DNA binding ability of MVI in the HeLaScribe lysate. This effect was observed in a concentration-dependent manner, consistent with our anisotropy data. The addition of 10 μM NDP52 did not decrease the transcription yield, suggesting that the Dab2 effect is not due to sequestering MVI through binding to the protein. The compromised transcription activity that we observed following antibody-depletion of MVI can be partially rescued through the addition of recombinant MVI (Figure 4d). Addition of recombinant MVI and NDP52 together can restore transcription to about 60 percent (Figure 4d), possibly due to the unfolding and dimerization of the protein. However, addition of recombinant MVI and Dab2 together failed to rescue the transcription yield, reinforcing that the observed effect is specific to Dab2, possibly through its interference with DNA binding. Taken together, our data suggest that Dab2 is a negative regulator of the transcription activity of MVI.

**Fig. 4.**
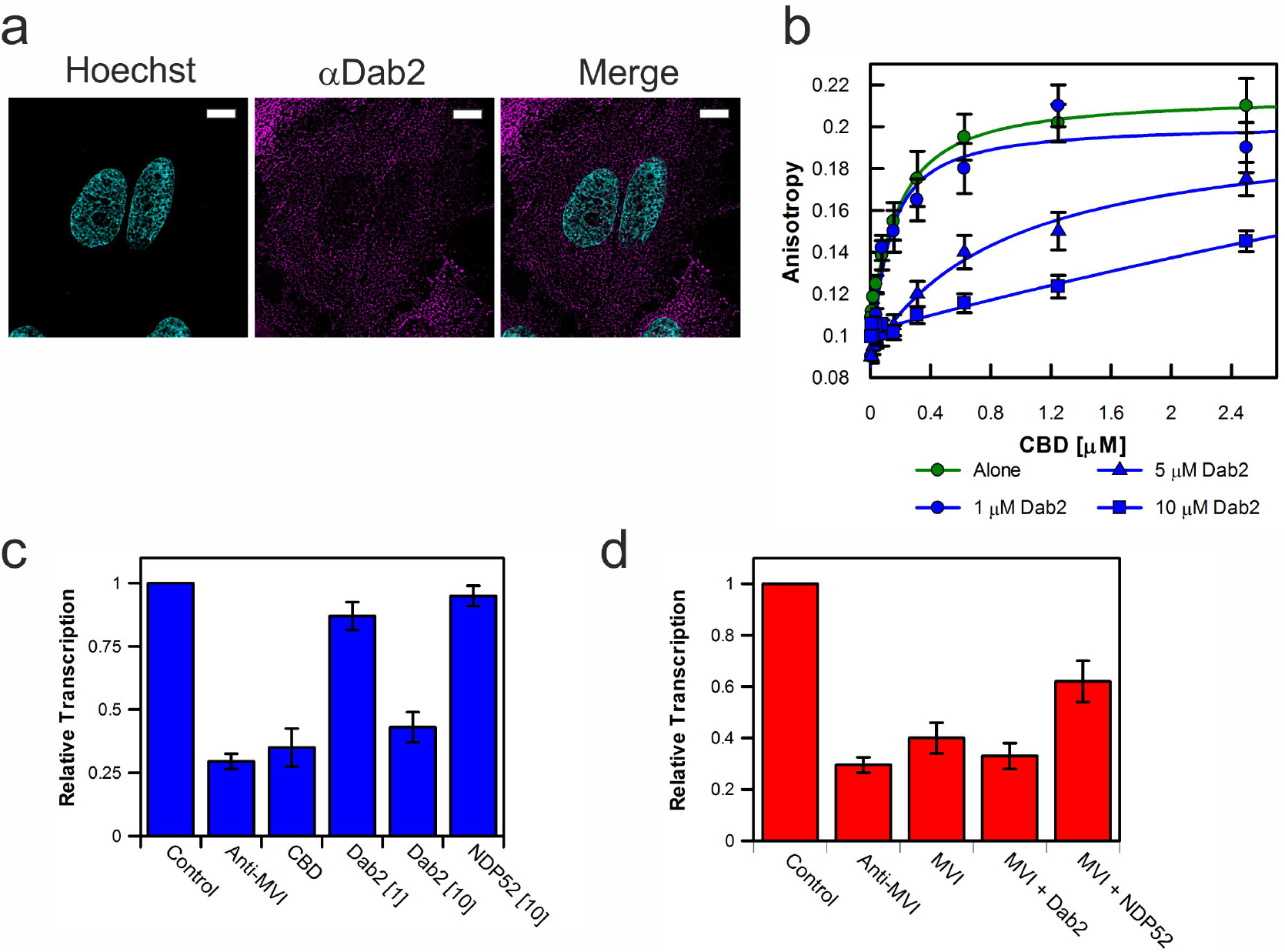
Dab2 represses the *in vitro* activity of nuclear myosin VI. (a) Immunofluorescence staining against Dab2 (magenta) combined with DNA staining (Cyan) in HeLa cells. Scale bar 10 μm. (b) Fluorescence anisotropy titrations of the CBD against a 40 bp fluorescein amidite (FAM)-DNA (50 nM) with the highlighted concentrations of Dab2. Data fitting was performed as described in Methods (*K*_d_ +/− SEM n = 3 independent experiments). (c) *In vitro* transcription by HelaScribe extracts following antibody depletion as described in Methods, or in the presence of CBD at 25 μM, Dab2 at 1 or 10 μM and NDP52 at 10 μM. Samples were normalized to a non-depleted control reaction (error bars represent SEM from 5 independent experiments). (d) *In vitro* transcription following antibody depletion and rescue using recombinant MVI (1 μM), NDP52 (10 μM) or Dab2 (10 μM), as described in Methods (Error bars represent SEM from 5 independent experiments).

### Effect of binding partner competition upon the nuclear functions of myosin VI

We have shown how two different binding partners can have contrasting effects on the biochemical properties and activity of MVI *in vitro*. On this basis, we then wanted to explore the impact of different binding partners on the cellular function of MVI, and in particular its nuclear role in breast cancer cell line MCF-7. MVI is also distributed throughout the cell body, including the nucleus (Figure 5a). In these cells, MVI has been already shown to be required for the expression of genes responsive to estrogen receptor (ER) signalling. When MCF-7 cells underwent siRNA knockdown of MVI, or treatment with TIP, the small molecule inhibitor of MVI, there was a decrease in the expression of estrogen-activated *PS2* and *GREB1* (Figure 5b). The impact of both treatments was similar, with a decrease of 70-80 percent for *PS2* and 30-40 percent for *GREB1*. Inhibition of MVI with TIP has already been shown to decrease the transcription yield *in vitro* (Cook et al., 2018). Based on our data, we can now directly link the motor activity of MVI to ER signalling. The genes that are under the control of the estrogen receptor relate, amongst others, to cell growth. Given the role of MVI in the expression of these genes, we then assessed the effect of MVI knockdown on MCF-7 growth (Figure 5c). Indeed, the loss of MVI attenuates cell growth, as it would be expected from the impact on gene expression. MCF-7 cells, along with many other estrogen receptor positive breast and ovarian cancer cell lines, have lost or attenuated Dab2 expression (Supplementary Figure 2a). Moreover, the reintroduction of Dab2 to these cell lines has been suggested to suppress tumourgenicity (He et al., 2001). Based upon the impact of MVI on the ER driven gene expression and our *in vitro* data demonstrating the impact of Dab2 upon MVI associated transcription, we wished to explore whether reintroduction of Dab2 in MCF-7 could lead to a perturbation of in the expression of ER target genes. To this end, we transiently expressed full length Dab2-mRFP in MCF-7 cells (Supplementary Figure 2a and b) and then monitored the expression of *PS2* and *GREB1*. Interestingly, the presence of Dab2 in MCF-7 resulted in a 20-30 percent decrease in the expression of these ER-target genes. To confirm that this decrease is due to the targeting of MVI, we transiently expressed in MCF-7 two truncations of Dab2 (Supplementary Figure 2c): the Dab2_649-770_ region which contains the MVI binding site and is the one used in our biochemical assays, and the Dab21-648 which cannot associate with MVI (Spudich et al., 2007). As expected, the Dab2_649-770_ resulted in a 40-50 percent decrease in the expression of the two ER target genes, whereas the Dab21-648 did not have any effect, confirming that the observed decrease in gene expression is indeed due to the interaction of Dab2 with MVI. Overall, our observations suggest that Dab2 can function in the cell as a negative regulator of MVI in the expression of ER-target genes.

**Fig. 5.**
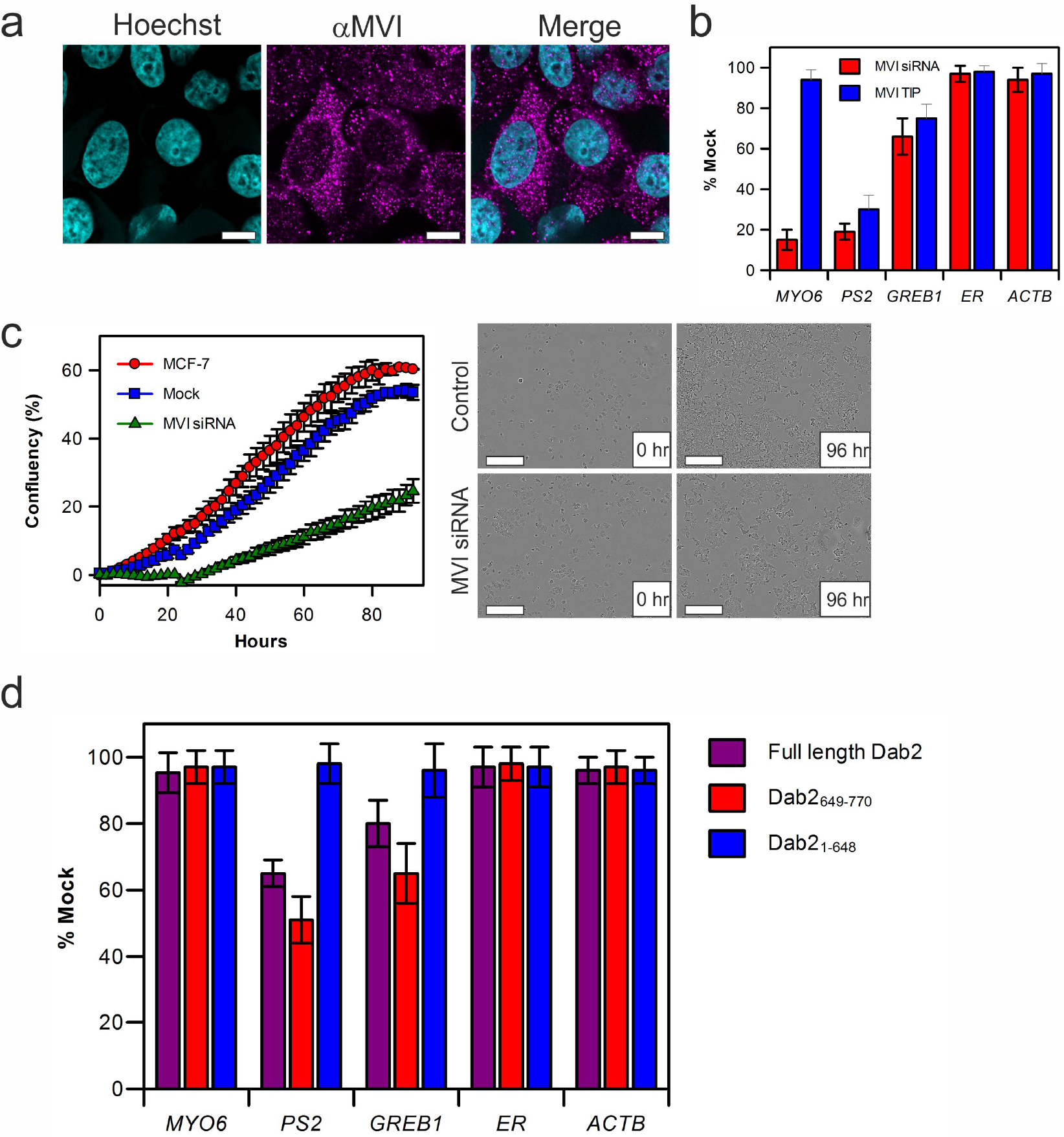
Inhibition of myosin VI in estrogen linked gene expression. (a) Immunofluorescence staining against MVI (magenta) combined with DNA staining (Cyan) in HeLa cells. Scale bar 10 μm. (b) Expression of estrogen receptor gene targets following siRNA knockdown of MVI (red), or TIP treatment (blue) in MCF-7 cells. Expression is plotted as a percentage of expression in mock cells. MYO6 reports on the success of the siRNA knockdown, whilst *ESR1* and *ACTB* were used to reflect global changes in transcription. (c) Real-time growth of MCF-7 cells (red) and corresponding measurements following MVI siRNA knockdown (green) and mock transfection control (blue). Data represent three independent measurements and error bars show SEM. Example images at start and end time points are shown. Scale bar 300 μm. Western-blot against Dab2 following transient transfection in to MCF-7 cells is shown in Supplementry Figure 2A. (d) Expression of estrogen receptor target genes following transient transfections of Dab2 (magenta), Dab21-648 (blue) and Dab2_649-770_ (red) in to MCF-7 cells. Expression is plotted as a percentage of expression in non-transfected cells. *MYO6*, *ESR1* and *ACTB* were used to reflect global changes in transcription (Error bars represent SEM from 3 independent experiments).

## Discussion

The structural regulation of MVI controls its biochemical properties and therefore directly impacts on the cellular function of this motor protein. Here, we have explored what is the impact of binding partners on the conformation of MVI, how these interactions are regulated and how this regulation varies between isoforms. We have then investigated how the selectivity of MVI for its binding partner can affect its bio-chemical properties and intracellular functions. We have revealed that the conformation of MVI and the structural rear-rangements following its association with its binding partners are independent of isoform and binding partner. Therefore, the structural regulation of MVI seems to follow a single general mechanism, which is summarised by the following model: MVI exists as a folded monomer which interacts with binding partners through the CBD. These interactions lead to MVI unfolding, and subsequent dimerization of the protein through its tail domain. As shown previously, this generates a processive motor protein (Fili et al., 2017). Moreover, here we have shown that this widely applied mechanism of structural regulation is under the control of a finely tuned inter-play between binding partners and the MVI isoforms. In the case of the LI isoform, structural studies have revealed how the RRL motif is blocked by the LI helix (Wollscheid et al., 2016), allowing only the WWY partners, such as Dab2, to bind to MVI. We have confirmed this here biochemically, by measuring the binding partner interactions at each site, using Dab2 as a representative example. In contrast, the NI isoform, in which both the RRL and WWY motif are readily accessible, should be able associate with any partner. We have therefore explored the mechanism underlying the partner selectivity in this case. We have revealed that the NI isoform can enact selectivity through the differential affinity between the two sites, with the RRL motif having stronger binding affinity over the WWY. In this way, the selectivity for binding partners is based upon the relative concentration of WWY versus RRL partners. Such a regulation could be achieved through changes in the local intracellular levels, or through a global change in expression levels. For instance, the loss of a WWY partner would perturb the dynamics leading to an enhanced role of RRL binding partners. This mechanism of selectivity is particularly relevant for the regulation of nuclear MVI, which is the NI isoform (Fili et al., 2017). Here, we have demonstrated the impact of a concentration driven interaction with the low affinity binding partner Dab2. Indeed, interaction with Dab2 blocked the ability of MVI to bind DNA and subsequently its transcription activity in vitro. The impact of Dab2 of the biochemical properties of MVI raised questions about its impact on the cellular functions of MVI. Dab2 has been proposed to function as a tumour suppressor, however, the underlying mechanism remains elusive. Here, we have proposed that at least part of this activity could relate to the down-regulation of nuclear MVI in transcription. As nuclear MVI is linked to estrogen receptor gene expression, this would in turn attenuate the activity of the estrogen receptor, subsequently leading to a decrease in tumorgenicity. Conversely, loss of a Dab2 would perturb the dynamics leading to an enhanced role of RRL binding partners, like NDP52 leading to enhanced MVI transcription activity. Cancer cell lines, such as the MCF-7, over-express the NI isoform of MVI, which is the one able to translocate to the nucleus, and they are therefore primed for transcriptional activity. This is further enhanced in MCF-7 cells by the loss of Dab2 expression, which relieves the Dab2-mediated negative regulation. Overall, this would lead to a high level of ER activity, which is MVI-dependent. Interestingly, reintroduction of Dab2 in these cells has an effect on their tumorigenic potential (He et al., 2001). In this study, we have confirmed that reintroduction of Dab2 perturbs the transcription landscape downstream the ER and revealed this is due to MVI targeting. We therefore suggest that the down-regulation of the transcriptional activity of MVI is indeed part of the role of Dab2 as a tumour suppressor. In summary, this study has allowed us to gain new insights into the regulation of MVI. We have shown that MVI exists as a folded monomer which is unfolded by binding partner, which can lead to its dimerization. Moreover, we have revealed that these conformation rearrangements are independent of the MVI isoform and binding partner. However, the mechanism of selectivity of binding partners is isoform depended. While the LI MVI employs a structural selection for its binding partners, the selectivity of the NI isoform is regulated by the relative expression levels of the partners. Finally, we provide an example of how the intracellular levels of MVI binding partners can modulate the cellular function of the protein. We propose that the tumour suppressor activity of Dab2 is, in part, related to the down-regulation of estrogen receptor target gene activation by nuclear MVI. These insights open new avenues for exploring how the activity of this multi-functional motor protein is regulated within the nucleus and the cytoplasm, as well.

## Methods

See Supplementary Information.

## Author Contributions

NF, YHG, B. Aston, AdS and CPT conceived the experiments. NF, YHG, AdS, REG B.Aston, B.Alamad and CPT performed experiments and analysed data. LW and MMF assisted with microscopy NF, YHG and CPT wrote the manuscript.

## ACKNOWLEDGEMENTS

We thank the MRC (MR/M020606/1) and STFC (19130001) for funding. We also thank Darren Griffin for sharing of equipment.

