## Supplementary Information for "Binding partner regulation of Myosin VI: Loss of tumour-suppressor Dab2 leads to enhanced activity of nuclear myosin"

### SUPPLEMENTARY FIGURES

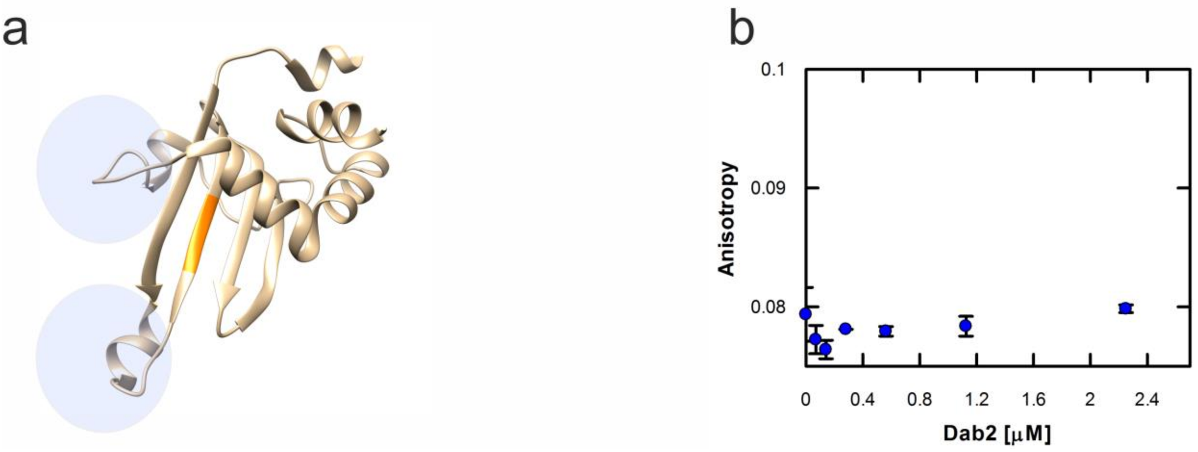


**Supplementary Figure 1 Characterisation of Dab2 DNA binding.**

(a) Structure of the C-terminal CBD with DNA binding sites highlighted (blue circles). The WWY motif which binds Dab2 is shown in orange. (PDB:2KIA(Yu et al., 2009)).

(b) Fluorescence anisotropy titrations of Dab2 against a 40 bp fluorescein amidite (FAM)-DNA (50 nM). Data fitting was performed as described in Methods ($K_d$ +/- SEM $n$ = 3 independent experiments).

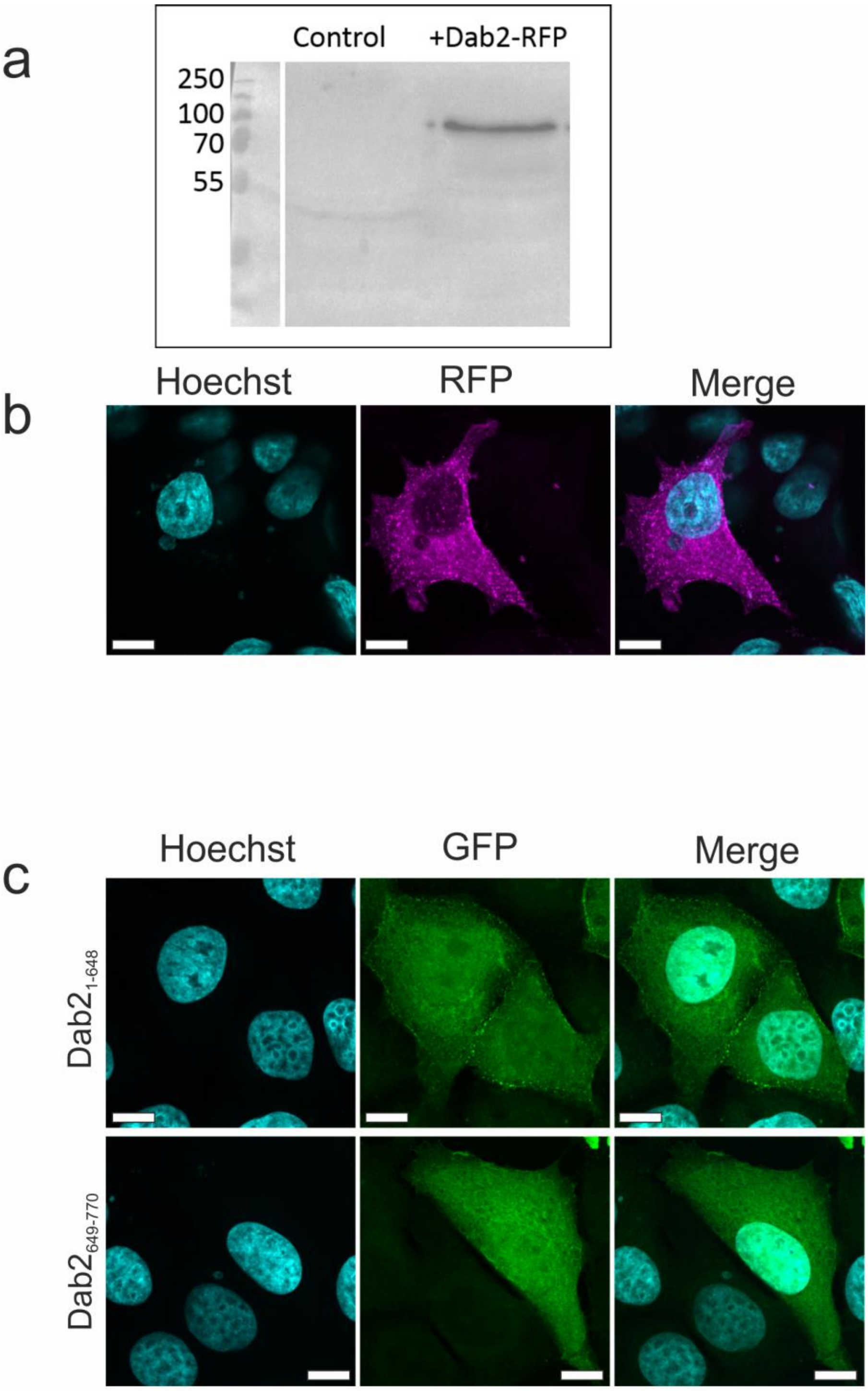


**Supplementary Figure 2 Transfection of Dab2 in MCF-7 cells.**

(a) Western Blot against Dab2 in MCF-7 cells (Control) and MCF-7 cells transiently transfected with Dab2-RFP.

(b) Representative images of transiently expressed Dab2-RFP in MCF-7 cells combined with DNA staining (Cyan).

(c) Representative images of transiently expressed GFP-Dab2$_{1\text{-}648}$ and GFP-Dab2$_{649\text{-}770}$ in MCF-7 cells combined with DNA staining (Cyan). Scale bar 10 μm in all images.

### METHODS

#### *Constructs*

A list of constructs and PCR primers are provided in Supplementary Table 1 and 2, respectively. Constructs generated in this work are described below: The following human full-length Myosin VI mutants (EGFP-C3-NI-MVI-RRL-AAA and EGFP-C3-NI-MVI-WWY-WLY) were a kind gift from F. Buss (CIMR). The full length human Dab2-mRFP was isolated by PCR from pET28a-Dab2-RFP plasmid, restriction digested with NheI and NotI and cloned into the pZsgreen1-N1 backbone following removal of Zsgreen1. The human Dab2 truncations were generated by PCR isolation of the Dab2 fragment containing the Myosin VI binding site (aa 649-770) and the rest of Dab2 (aa 1-648) which does not contain the Myosin VI binding site from the above Dab2-mRFP plasmid. The PCRs were restriction digested by XhoI and SacII and cloned into pEGFP-C3.

#### *Protein expression and purification in Escherichia coli*

Recombinant constructs were expressed in *E.coli* BL21 DE3 cells (Invitrogen) in Luria Bertani media. Proteins were purified by affinity chromatography (HisTrap FF, GE Healthcare). The purest fractions were further purified through a Superdex 200 16/600 column (GE Healthcare).

#### *Protein Expression using Baculovirus system*

Full-length myosin VI NI and LI and *Xenopus* calmodulin were expressed in *Sf9* and *Sf21* (*Spodoptera frugiperda*) insect cells using the Baculovirus expression system. *Sf9* cells were cultured in suspension in sf900 media (Gibco) at 27ºC to generate the P1-3 recombinant baculovirus stocks. Finally, expression of recombinant proteins was set up by infecting *sf21* cells with the P3 viral stock in ExCell 420 media (Sigma). The cells were harvested by centrifugation (as above) for protein purification after 4 days. Prior to sonication, an additional 5 mg Calmodulin was added with 2 mM DTT. After sonication, 5 mM ATP and 10 mM $MgCl_2$ were added and the solution was rotated at 4 °C for 30 min before centrifugation (20,000g, 4°C, 30 min). Then, the cell lysate was subjected to the purification steps described above.

***Protein labelling***

Proteins were transferred into 50 mM Na-phosphate (pH 6.5) using a PD10 desalting column. Samples were then incubated with a 5-fold excess of dye for 4 hours, rotating at 4°C**.** Excess dye was removed using a PD10 desalting column pre-equilibrated with 50 mM Na-Phosphate, 150 mM NaCl and 1 mM DTT. Labelling efficiency was calculated based on the absorbance at 280 nm and the absorbance maximum of the dye. Typical efficiency was 90%, whereby the less than complete labelling was taken as an indicator for a single dye per protein. This was tested for isolated preparations in mass spectroscopy, which revealed both an unlabelled and single labelled population.

***Cell culture and Transfection***

HeLa (ECACC 93021013) and MCF7 (ECACC 86012803) cells were cultured at 37ºC and 5% $CO_2$, in Gibco MEM Alpha medium with GlutaMAX (no nucleosides), supplemented with 10% Fetal Bovine Serum (Gibco), 100 units/ml penicillin and 100 µg/ml streptomycin (Gibco). For the transient expression of MVI isoforms and mutants, full length Dab2 and truncations, HeLa cells and/or MCF7 cells grown on glass coverslips were transfected with EGFP-NI-MVI, EGFP-LI-MVI, EGFP-NI-MVI-RRL-AAA, EGFP-NI-MVI-WWY-WLY, Dab2-mRFP, EGFP-Dab2-1-648 and EGFP-Dab2-649-770 constructs using Lipofectamine 2000 (Invitrogen), following the manufacturer's instructions. Depending on the construct, 24 h - 72 h after transfection, cells were subjected to nuclear staining using Hoechst 33342 (Thermo Scientific), fixed and analysed or subjected to indirect immunofluorescence (see below).

Full length Dab2-RFP cDNA was electroporated into MCF7 cells using BioRad Gene Pulser Xcell TM electroporation system. After trypsinization, harvested cells were washed with 1X PBS, counted and $1.5\times10^6$ cells were resuspended in 800ul of cold Opti-MEM media and the cell suspension was kept on ice. 10ug of DNA was added to the cell suspension and the mixture was transferred to the Biorad 4mm cuvette. The cells were then pulsed using exponential-decay protocol with a voltage of 300V and a capacitance of 350µF. Cells were allowed to recover for 5 minutes after which warm complete media (MEM) was added to them and they were plated in a 6 well plate at a density of $0.5\times10^6$ cells per well. The cells were collected for protein or RNA extraction after 48 hours.

***Immunofluorescence***

Transfected and non-transfected HeLa or MCF-7 cells were fixed for 15 min at room temperature in 4% (w/v) paraformaldehyde (PFA) and residual PFA was quenched for 15 min with 50 mM ammonium chloride. All subsequent steps were performed at room temperature. Cells were permeabilised and simultaneously blocked for 15 min with

0.1 % (v/v) Triton X-100 and 2 % (w/v) BSA in PBS. Cells were then immuno-stained against the endogenous proteins by 1 hour incubation with the indicated primary and subsequently the appropriate fluorophore-conjugated secondary antibody (details below), both diluted in 2% (w/v) BSA in PBS. The following antibodies were used at the indicated dilutions: Rabbit anti-MVI (1:200, Atlas-Sigma HPA0354863-100UL), Mouse anti-Dab2 (1:100, Abcam ab88590) ,donkey anti-mouse Alexa Fluor 488-conjugated (1:500, Abcam Ab181289), donkey anti-mouse Alexa Fluor 555-conjugated (1:500, Abcam Ab150110), donkey anti-rabbit Alexa Fluor 488-conjugated antibody (1:500, Abcam Ab181346) and donkey anti-rabbit Alexa Fluor 555-conjugated antibody (1:500, Abcam Ab150074). Coverslips were mounted on microscope slides with Mowiol (10% (w/v) Mowiol 4-88, 25% (w/v) glycerol, 0.2 M Tris-HCl, pH 8.5), supplemented with 2.5% (w/v) of the anti-fading reagent DABCO (Sigma).

For co-localisation analysis, ROIs were drawn around individual cells (minimum 10 cells and 5 stacks per cell). Pearson's coefficients were obtained with the JACoP plugin(Bolte and Cordelieres, 2006) for ImageJ.

***Immunoblot Analysis***

The total protein concentration was determined by Bradford Assay (Sigma) following the manufacturer’s instructions. Cell lysates were heat-denatured and resolved by SDS-PAGE. The membrane was probed against the endogenous proteins by incubation with mouse Anti-Dab2 polyclonal antibody (1:1000, Abcam ab88590) and subsequently a goat anti-mouse antibody coupled to horseradish peroxidase (1:15000, Abcamab97023). The bands were visualised using the ECL Western

Blotting Detection Reagents (Invitrogen) and the images were taken using Syngene GBox system. Images were processed in ImageJ.

***Fluorescence Imaging***

Cells were visualised using either the ZEISS LSM 880 confocal microscope or the widefield Olympus IX71 microscope. The former was equipped with a Plan-Apochromat 63x 1.4 NA oil immersion lens (Carl Zeiss, 420782-9900-000). Three laser lines, i.e. 405 nm, 488 nm and 561 nm, were used to excite the fluorophores, i.e. Hoechst, GFP and RFP, respectively. The built-in dichroic mirrors (Carl Zeiss, MBS-405, MBS-488 and MBS-561) were used to reflect the excitation laser beams on to cell samples. The emission spectral bands for fluorescence collection were 410 nm-524 nm (Hoechst), 493 nm-578 nm (GFP) and 564 nm-697 nm (RFP). The detectors consisted of two multi anode photomultiplier tubes (MA-PMT) and 1 gallium arsenide phosphide (GaAsP) detector. The green channel (GFP) was imaged using GaAsP detector, while the blue (Hoechst) and red (RFP) channels were imaged using MA-PMTs. ZEN software (Carl Zeiss, ZEN 2.3) was used to acquire and render the confocal images. The later was equipped with an PlanApo 100xOTIRFM-SP 1.49 NA lens mounted on a PIFOC z-axis focus drive (Physik Instrumente, Karlsruhe, Germany), and illuminated with an automated 300W Xenon light source (Sutter, Novato, CA) with appropriate filters (Chroma, Bellows Falls, VT). Images were acquired using a QuantEM (Photometrics) EMCCD camera, controlled by the Metamorph software (Molecular Devices). The whole volume of cells was imaged by acquiring images at z-steps of 200 nm. Widefield images were deconvolved with the Huygens Essential version 17.10 software. Confocal Images were deconvolved using the Zeiss Zen2.3 Blue software, using the regularised inverse filter method. All images were then analysed by ImageJ.

***Incucyte***

Cells were seeded onto 96-well tissue culture dishes at equal densities in 6 replicates. After attachment over-night, cells were transfected with MVI siRNA. Photomicrographs were taken every hour using an IncuCyte live cell imager (Essen Biosciences, Ann Harbor, MI) and confluency of cultures was measured using IncuCyte software. Confluency values between wells were normalised to initial confluency for comparison.

***RNA extraction and RT-qPCR***

RNA from Dab2-RFP transfected or non-transfected MCF7 cells was extracted using Gene Jet RNA purification kit (Thermo scientific) according to manufacturer's protocol. The RNA concentration was measured using Geneflow Nanophotometer and RT-qPCR was performed with one-step QuantiFast SYBR Green qPCR kit (Qiagen) using 50ng of RNA in each sample. A list of qPCR primers is given in Supplementary Table 3.

***DNA Substrates***

DNA substrate ds40 consisted of Labeled (TTAGTTGTTCGTAGTGCTCGTCTGGCTCTGGATTACCCGC*FAM)
and unlabelled (GCGGGTAATCCAGAGCCAGACGAGCACTACGAACAACTAA)
oligonucleotides purchased from IDT. To form duplex DNA substrates, oligonucleotides were mixed at equimolar concentrations at either 50 µM in water or a buffer containing 50 mM Tris.HCl at pH 7.5, 150 mM NaCl, and 3 mM MgCl2.

***In vitro transcription***

The DNA template was the pEGFP-C3 linearized plasmid containing the CMV promoter which would generate a 130-base run-off transcript. The HelaScribe (Promega) reactions were performed in triplicates, through two independent experiments, according to the manufacturer's instructions. The reactions were performed for 60 min at 25°C.

Reactions were also performed following pre-clearance with the MVI antibody. Protein G Dynabeads (Invitrogen) were prepared according to manufacturer's instructions before being loaded with 4 μg antibody. Samples were incubated for 30 min on ice and beads were extracted immediately before performing the transcription reaction.

For quantification, mRNA was purified using Gene Jet RNA purification kit (Thermo scientific) according to manufacturer's protocol and RT-qPCR was performed with one-step QuantiFast SYBR Green qPCR kit (Qiagen).

***Steady-state ATPase Activity of MVI***

$Ca^{2+}$-actin monomers were converted to $Mg^{2+}$-actin with 0.2 mM EGTA and 50 μM $MgCl_2$ before polymerizing by dialysis into 20 mM Tris.HCl (pH7.5), 20 mM imidazole (pH 7.4), 25 mM NaCl and 1 mM DTT. A 1.1 molar equivalent of phalloidin (Sigma) was used to stabilize actin filaments.

Steady-state ATPase activities were measured at 25 °C in KMg50 buffer (50 mM KCl, 1 mM MgCl2, 1 mM EGTA, 1 mM DTT, and 10 mM imidazole, pH 7.0). Supplemented with the NADH-coupled assay components, 0.2 mM NADH, 2 mM phosphoenolpyruvate, 3.3 U $ml^{-1}$ lactate dehydrogenase, 2.3 U $ml^{-1}$ pyruvate kinase and various actin concentrations (0 – 30 μM). The final [Mg.ATP] was 5 mM and MVI concentration was 100–300 nM. The assay was started by the addition of MVI. The change in absorption at $OD_{340}$ nm was followed for 5 min. The $k_{cat}$ and $K_{actin}$ values were determined by fitting the data to equation 1.

$$Rate = V_o + \left(\frac{k_{cat}[Actin]}{K_{actin}+[Actin]}\right)$$

$V_o$ is the basal ATPase activity of MVI, $k_{cat}$ is the maximum actin-activated ATPase rate and $K_{actin}$ is the concentration of actin needed to reach half maximal ATPase activity.

#### *Titration measurements*

All reactions were performed at 25 °C in a buffer containing 50 mM Tris·HCl (pH 7.5), 150 mM sodium chloride and 1 mM DTT in a final volume of 100 μL. Measurements were performed using a ClarioStar Plate Reader (BMG Labtech).

Intensity measurements were performed at the following wavelengths: FITC (ex. 490nm), Alexa Fluor 555 (ex. 555nm). FITC to Alexa Fluor 555 FRET measurements were performed using the following wavelengths ex. 470nm and em. 575nm. Anisotropy was measured with the instrument in the T format, allowing simultaneous acquisition of parallel ($I_{//}$) and perpendicular ($I_{\perp}$) components using BMG filter-sets for fluorescein (Ex. 482/16-10, Dichroic LP504 and em. 530/-40).

#### *Analysis of kinetic data*

For Fluorescence Anisotropy titrations: Anisotropy was calculated, as described below, based upon established procedures (Brownbridge et al., 1993)'(Soh et al., 2015; Toseland, 2014; Toseland and Geeves, 2014).

Total fluorescence intensity ($F_t$) is given by:

$$F_t = \sum c_i F_i$$

Total anisotropy ($A_t$) is given by:

$$A_t = \frac{\sum c_i F_i A_i}{F_t}$$

Where $c_i$ is the concentration of species *i*, $F_i$ is the fluorescence intensity per unit of concentration and $A_i$ is the anisotropy. This is calculated from the parallel and perpendicular fluorescence intensity (*I*) in relation to the plane of excitation by:

$$A_i = \frac{I_{parallel} - I_{perpendicular}}{I_{parallel} + 2I_{perpendicular}}$$

As anisotropy is additive for multiple fluorescence species in solution, it is used to give a measure of their relative concentrations. For MVI (and various constructs) there are two fluorescence species, DNA and MVI.DNA. The total anisotropy can then be calculated in terms of the dissociation constant ($K_d$) for the MVI.DNA complex:

$$A_t = \frac{A_{DNA}([DNA]_t - [MVI.DNA]) + A_{MVI.DNA}Q[MVI.DNA]}{[DNA]_t - [MVI.DNA] + Q[MVI.DNA]}$$

Where

$$[MVI.DNA] = \frac{([MVI]_t + [DNA]_t + K_d) - \sqrt{([MVI]_t + [DNA]_t + K_d)^2 - 4[MVI]_t[DNA]_t}}{2}$$

And where $[MVI]_t$ and $[DNA]_t$ are the total concentrations for each reactant. [MVI.DNA] is the concentration of the protein-bound DNA complex. Q is the fluorescence intensity of MVI.DNA relative to DNA. The anisotropy data were fitted to obtain dissociation constants based on the above equations using GraFit fitting software (Leatherbarrow, R. J. (2001) Grafit Version 5, Erithacus Software Ltd., Horley, U.K.).

For the FRET titrations: The 575 nm intensity data was corrected for the increase in intensity due to a small direct excitation. This background signal was subtracted from the dataset to leave the FRET values. The titration curves for the $MVI_{TAIL}$ interactions were fitting to a binding quadratic equation:

$$[Complex] = \frac{([FITC]_t + [AF555]_t + K_d) - \sqrt{([FITC]_t + [AF555]_t + K_d)^2 - 4[FITC]_t[AF555]_t}}{2}$$

***Data Availability***

The data supporting the findings of this study are available from the corresponding author on request.

**Supplementary Table 1. Recombinant DNA.**

| **Construct (Residue numbers)** | **Source** |
|---|---|
| Human pEGFP-C3 myosin VI (large insert) (1-1284) | F. Buss (CIMR) |
| Human pEGFP-C3 myosin VI (non insert) (1-1253) | F. Buss (CIMR) |
| Human pFastBacHTB GFP NI myosin VI | F. Buss (CIMR) |
| Human pFastBacHTB GFP LI myosin VI | F. Buss (CIMR) |
| Xenopus pFastBac1 Calmodulin (1-end) | J. Sellers (NIH) |
| Human pET151 NDP52 (1-end) | Ref Fili et al 2017 |
| Human pET151 EGFP-$MVI_{TAIL}$-RFP (814-1253) | Ref Fili et al 2017 |
| Human pET151 $MVI_{TAIL}$ 814-1253 | Ref Fili et al 2017 |
| Human pET151 $_{N}MVI_{TAIL}$ 814-1060 | Ref Fili et al 2017 |
| Human pET28 CBD 1060-1253 | Ref Fili et al 2017 |
| Human pET28 $MVI_{TAIL}$ (large insert) (814-1284) | Ref Fili et al 2017 |
| Human pET28 CBD (large insert) (1037-1284) | Ref Fili et al 2017 |
| Human pFastbacHTB NI MVI (1-1253) | Ref Fili et al 2017 |
| Xenopus pET28 Calmodulin | Ref Fili et al 2017 |
| Human pET151 Dab2 (649-770) | Synthetic Gene – This Study |
| Human pZsgreen1 backbone Dab2 RFP | This Study |
| Human pEGFP-C3 myosin VI (non insert) WWY-WLY | This Study |
| Human pEGFP-C3 myosin VI (non insert) RRL-AAA | This Study |
| Human pEGFP-C3 Dab2 (1-648) | This Study |
| Human pEGFP-C3 Dab2 (649-770) | This Study |
| Human pET28 Dab2 RFP | F. Buss (CIMR) |

**Supplementary Table 2. PCR Primers.**

| Sequence | Use |
|---|---|
| AGCTAGCTGCATGGCTGACCAACTG | pET28 Calmodulin For |
| TTTTGCGGCCGCTCACTTTGCTGTCATC | pET28 Calmodulin Rev |
| CTAGGCGGCCGCCCGAGGATGGAAAGCCCGTTTG | pFastbacHTB NI Myosin VI |
| TTTTCTCGAGTTATTTCAACAGGTTCTGCAGCATG | pFastbacHTB NI Myosin VI |
| CACC GAAGCCTGCATTAAAATGC | MVI$_{TAIL}$ 814-1253 For |
| CTATTTCAACAGGTTCTGCAGCAT | MVI$_{TAIL}$ 814-1253 Rev |
| CTAGGCGGCCGCCCGAGGATGGAAAGCCCGTTTG | pFastBacHTB NI MVI |
| TTTTCTCGAGCTAGCATTTTAATGCAGGCTTC | pFastBacHTB NI MVI |
| CTAGGCGGCCGCCCGAGGATGGAAAGCCCGTTTG | pFastBacHTB NI MVI |
| TTTTCTCGAGCTAGATGGTATCACGTAGTTCTGC | pFastBacHTB NI MVI |
| AGCTAGCGCAGAACTACGTGATACCATC | CBD For |
| TTTTGCGGCCGCTATTTCAACAGGTTCTGCAGCAT | CBD Rev |
| AGCTAGCGCAGAACTACGTGATACCATC | $_{N}$CBD For |
| TTTTGCGGCCGCTATGCTGGGTTTTGCTGAGG | $_{N}$CBD Rev |
| AGCTAGCGCTCAGATTCCTGCCAGG | $_{C}$CBD For |
| TTTTGCGGCCGCTATTTCAACAGGTTCTGCAGCAT | $_{C}$CBD Rev |
| AGCTAGCGGGGCAGAACTCAGCACTG | CBD Plus LI For |
| TTTTGCGGCCGCTATTTCAACAGGTTCTGCAGCAT | CBD Plus LI Rev |
| CACC GAAGCCTGCATTAAAATGC | $_{N}$MVI$_{TAIL}$ For 814-1060 |
| CTAGATGGTATCACGTAGTTCTGC | $_{N}$MVI$_{TAIL}$ Rev 814-1060 |
| AGCTAGCGCCGAACTGCGTGATACCA | CBD W1221A For |

| | |
|---|---|
| TTTTGCGGCCGCTATTATTTCAGCAGGCTCTGCAGC | CBD W1221A Rev |
| CTTGCAGAGAAGAATTTCATGCGGCAGCAAAAGTGTATCATGC | RRL/AAA For |
| GATTTCCAAGCATGATACACTTTTGCTGCCGCATGAAATTCTTC | RRL/AAA Rev |
| CTTGCAGAGAAGAATTTCATAGGAGAGCAAAAGTGTATCATGC | RRL/RRA For (L1118A) |
| GATTTCCAAGCATGATACACTTTTGCTCTCCTATGAAATTCTTC | RRL/RRA Rev (L1118A) |
| CTTGCAGAGAAGAATTTCATGCGAGACTAAAAGTGTATCATGC | RRL/ARL For (R1116A) |
| GATTTCCAAGCATGATACACTTTTAGTCTCGCATGAAATTCTTC | RRL/ARL Rev (R1116A) |
| GCAGAGAAGAATTTCATAGGGCACTAAAAGTGTATCATGCTTG | RRL/RAL Rev (R1117A) |
| CCAAGCATGATACACTTTTAGTGCCCTATGAAATTCTTCTCTGC | RRL/RAL Rev (R1117A) |
| GACCCTCAGAGTGCGGCAGCAGGCTGGTGGTATGC | SKKK/SAAA For |
| GCATACCACCAGCCTGCTGCCGCACTCTGAGGGTC | SKKK/SAAA Rev |
| GACTGGCCTGACTCGGAAGCGTGGTGCTGAGATCTTG | TRKR/TAAA For |
| CAAGATCTCAGCACCACGCTTCCGAGTCAGGCCAGTC | TRKR/TAAA Rev |
| GCCATGGCGCAGAACGCGGCGAAATAAGCCGAAG | LQSLL/AQSAA For |
| CTTCGGCTTATTTCGCCGCGTTCTGCGCCATGGC | LQSLL/AQSAA Rev |
| GATCGATAGTACATAAGGATTTCCTTACGCG | 500bp 5'Bio-Teg |
| CCAATTTTCGTTTGTTGAACTAATGGGTGC | 500 bp Rev |

**Supplementary Table 3. Primers for qPCR.**

| **Sequence** | **Use** |
|---|---|
| CATGGAGAACAAGGTGATCTG | TFF1/PS2 qPCR For |
| CACTGTACACGTCTCTGTCTG | TFF1/PS2 qPCR Rev |
| ATGGGAAATTCTTACGCTGGAC | GREB1 qPCR For |
| CACTCGGCTACCACCTTCT | GREB1 qPCR Rev |
| AGAGCTACGAGCTGCCTGAC | Human B-Actin qPCR For |
| AGCACTGTGTTGGCGTACAG | Human B-Actin qPCR Rev |
| AAGCTTCGATGATGGGCTTA | ESR1 qPCR For |
| AGGTGGACCTGATCATGGAG | ESR1 qPCR Rev |
| CCGAGCTCATCAGTGATGAGGC | Myosin VI qPCR For |
| CCAAGCATGATACACTTTTAGTCTCC | Myosin VI qPCR Rev |
| AAGGGCATCGACTTCAAGGA | CMV GFP RT-qPCR For In vitro Transcription |

| | |
|---|---|
| GGCGGATCTTGAAGTTCACC | CMV GFP RT-qPCR Rev In vitro Transcription |
